# The Adaptor Protein Complex 1 limits E-cadherin endocytosis during epithelial morphogenesis

**DOI:** 10.1101/2020.10.14.340372

**Authors:** Miguel Ramírez Moreno, Katy Boswell, Natalia A. Bulgakova

## Abstract

Intracellular trafficking regulates the distribution of transmembrane proteins including the key determinants of epithelial polarity and adhesion. The Adaptor Protein 1 (AP-1) complex is the key regulator of vesicle sorting, which binds a large number of specific cargos. We examined roles of the AP-1 complex in epithelial morphogenesis, using the *Drosophila* wing as a paradigm. We found that AP-1 knockdown leads to ectopic folds caused by trafficking defects of integrins. This occurs concurrently with an increase in the apical cell area and induction of cell death due to defects in E-cadherin trafficking. We discovered a distinct pool of AP-1 localizes at the apical Adherens Junctions, where it limits internalization of E-cadherin from the cell surface. Upon AP-1 knockdown, the accompanying hyperinternalization of E-cadherin induces cell death by an uncharacterised mechanism with a potential tumour-suppressive role. Simultaneously, cells increase expression of E-cadherin in a compensatory mechanism to maintain cell-cell adhesion.

## Introduction

Epithelial morphogenesis comprises the series of processes, such as tissue growth and deformation, that give origin to complex structures like organs from simpler epithelial sheets (Schock & Perrimon, 2002). These processes are driven by changes in properties of the participating cells, often regulated by transmembrane proteins at their surfaces (Schock & Perrimon, 2002; Heisenberg & Bellaïche, 2013). In particular, adhesion molecules constitute the cornerstone of interactions between individual cells and with their environment, facilitating morphogenesis (Halbleib & Nelson, 2006; Gumbiner, 1996). Cell-cell adhesions are mediated by proteins such as E-cadherin (E-cad), forming homophilic interactions at the apical Adherens Junctions (AJs) (Harris & Peifer, 2004; Takeichi, 1977). E-cad is subjected to a complex and dynamic regulation, enabling its contribution to numerous developmental processes and diseases such as cancer (Bruser & Bogdan, 2017; Halbleib & Nelson, 2006; Janiszewska *et al*, 2020). Another type of cell adhesion molecules, the integrins, anchor cells to the basal extracellular matrix (Domínguez-Giménez *et al*, 2007). By binding to basal ligands, integrins carry roles in both cell architecture and signalling during morphogenesis (Yamada & Miyamoto, 1995; Domínguez-Giménez *et al*, 2007; Lee & Streuli, 2014). Additionally, altered integrin adhesion changes basal tension, leading to ectopic folding of the tissue (Sui *et al*, 2018). However, how cell surface presentation of adhesion proteins is regulated and impacts of this regulation on tissue morphogenesis remain the biggest questions in the fields of cell and developmental biology (Kowalczyk & Nanes, 2012).

The major way to regulate adhesion protein presentation at the cell surface is via intracellular trafficking, whereby vesicles containing cargo proteins move across organelles and specific routes (Herrmann & Spang, 2015; Grant & Donaldson, 2009). Transported cargos are sent from and to numerous organelles, with the Trans Golgi Network (TGN) serving as the main sorting centre in the cell (Grant & Donaldson, 2009; Tan *et al*, 2019). Upon fate decision in the TGN, the endosomal machinery directs the cargo vesicles either to the plasma membrane through the Recycling Endosomes (REs) or towards lysosomes (Grant & Donaldson, 2009). Most of what is known about trafficking pathways comes from studies in single cells, which maximize data output about molecular details but do not inform about their functions in tissue contexts during development (Fölsch *et al*, 2001; Huang *et al*, 2019; Guo *et al*, 2012; Bruser & Bogdan, 2017; York *et al*, 2020).

One of the key regulator of intracellular trafficking is the Adaptor Protein Complex 1 (AP-1), which shuttles vesicles within the TGN/REs continuum (Grant & Donaldson, 2009; Bonifacino, 2014; Tan *et al*, 2019; Bonifacino & Rojas, 2006). In mammals, two isoforms of the AP-1 complex exist depending on the participating μ subunit: the ubiquitously expressed AP-1A and the tissue-specific AP-1B (Hase *et al*, 2013; Gravotta *et al*, 2007, 2012). Whereas AP-1A sorts basolateral cargos at the TGN and depends on activity on the small GTPase Arf1; AP-1B sorts proteins from REs to the basolateral membrane and seems to be specifically regulated by the small GTPase Arf6 (Fölsch, 2015; Shteyn *et al*, 2011; Ren *et al*, 2013). Intriguingly, AP-1B is also found at integrin-mediated focal adhesions, and although its function there is unclear, its loss correlates with highly migratory behaviour of metastatic cancer cells (Kell *et al.,* 2020). Two AP-1s are also found in *Caenorhabditis elegans,* where they restrict the basolateral location of E-cad (Shafaq-Zadah *et al*, 2012; Gillard *et al*, 2015). In the fruit fly, *Drosophila melanogaster*, a single AP-1 complex localizes to both TGN and REs, and is required for the maintenance of E-cad at the highly specialized ring canals during oogenesis (Loyer *et al*, 2015; Benhra *et al*, 2011; Burgess *et al*, 2011). At the same time, in specialized epithelia – the retina and sensory organs – it has only been linked to distribution of Notch pathway components and has no reported effects on apical-basal cell polarity or adhesion (Loyer *et al*, 2015; Benhra *et al*, 2011; Kametaka *et al*, 2012).

In this work, we demonstrate that the *Drosophila* AP-1 complex is necessary for the correct epithelial morphogenesis and architecture using the developing wing as a paradigm. There, the AP-1 complex regulates multiple aspects of morphogenesis including cell size and number, as well as tissue folding. This regulation is achieved by controlling the transport of adhesion proteins: E-cad and integrins. We have identified a subapical fraction of the AP-1 complex outside its canonical localization at the TGN/REs, and determined that the AP-1 complex regulates the endocytosis of E-cad from the plasma membrane. The excessive internalization of the E-cad following AP-1 knockdown triggers cell death that may act as a tumour suppressing mechanism. The surface levels of E-cad remain however stable due to the adjustment on its expression. Altogether, our results demonstrate the versatility of functions of the AP-1 complex in intracellular trafficking *in vivo,* and link the intracellular trafficking to tissue development and pathology, regulating cell shape and survival.

## Results and Discussion

We knocked down AP-1 using interfering RNA (RNAi) with the GAL4/UAS system (Brand & Perrimon, 1993). We employed the *engrailed* promoter (*en::* GAL4) to express GAL4 throughout development of wing primordia (imaginal discs) in a broad domain: their posterior compartments (Fig 1A-C). Their anterior compartments then served as internal controls with identical genotypes. Downregulation of AP-1μ or AP-1γ (Fig EV1A) – two out of four subunits of the AP-1 complex (Tan *et al*, 2019; Robinson, 2004) – led to defective morphology of adult wings, ranging from completely absent to severely disrupted posterior compartments (Fig 1A). A similar though weaker adult phenotype was caused by using *MSIO96::GAL4* driver (Fig EV1B), which expresses GAL4 throughout the wing pouch (presumptive region of the adult wing, Fig 1C).

**Figure 1.**
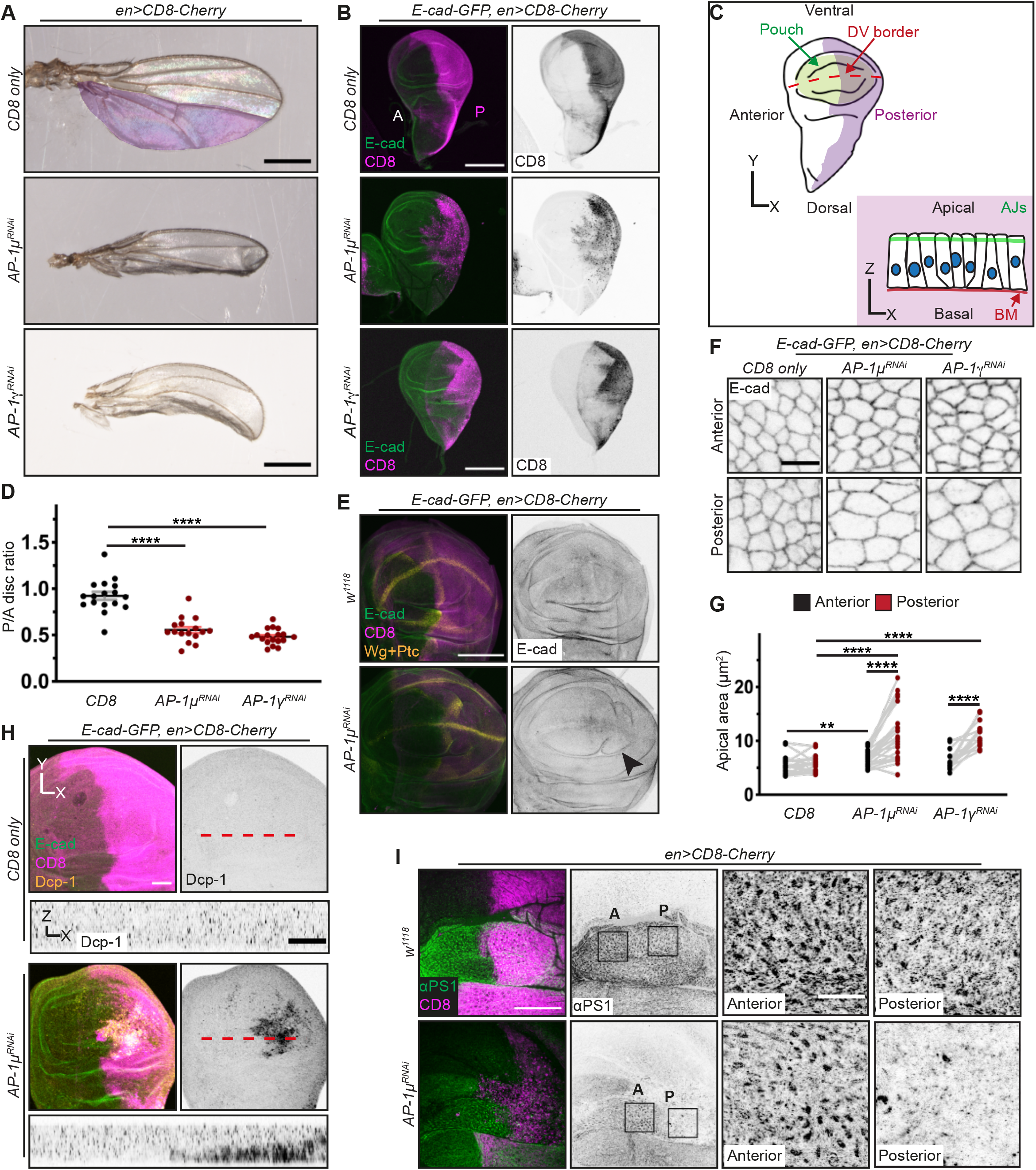
AP-1 regulates tissue development, cell morphology and cell survival in the wing disc. **A** Adult female fly wings expressing CD8-Cherry protein alone (top), with AP-1μ RNAi (middle), or with AP-1γ RNAi (bottom) in the posterior compartment. Scale bar: 0.5 mm. **B** Third instar larval wing discs expressing CD8-Cherry (magenta, left; inverted grayscale, right) alone (top), with AP-1μ RNAi (middle), or with AP-1γ RNAi (bottom) in the posterior compartment and showing E-cad-GFP (green, left). Letters indicate the Anterior (A) and Posterior (P, expressing CD8-Cherry) compartments in the wing disc Scale bar: 150 μm. **C** Top – cartoon of a third instar wing imaginal disc (top view) with highlighted posterior compartment (magenta), wing pouch (green) and dorsoventral border (DV, dashed red line). Bottom – a sagittal view of the pseudostratified epithelium in the pouch region with the apical Adherens Junctions (AJs, green) and the basal membrane (BM, red). **D** Posterior:Anterior (P/A) area ratios of the control discs and those expressing AP-1 RNAis. Dots represent individual discs (n=17, 16 and 18). ****P<0.0001 (Brown-Forsythe and Welch ANOVA test). **E** Wing pouch regions of discs expressing CD8-Cherry protein alone (top) or with AP-1μ RNAi (bottom) and co-stained with Wingless and Patched (Wg and Ptc, both in yellow). E-cad-GFP visualizes the tissue architecture (green, left; inverted grayscale, right). Arrowhead indicates an ectopic fold. Scale bar: 100 μm. **F** Apical view of the anterior (top) and posterior (bottom) compartments of discs expressing CD8-Cherry protein alone (left), with AP-1μ RNAi (middle), or AP-1γ RNAi (right). Cell outlines are visualized with E-cad-GFP (inverted grayscale). Scale bar: 5 μm. **G** The cell apical area of wing discs depicted in F, with each dot representing the cell average of an individual disc (n=15-23 discs/genotype). **P<0.01 and ****P<0.0001 (Wilcoxon and paired t-test, anterior versus posterior; and Kluskal-Walls or two-Way ANOVA tests, comparisons between genotypes). **H** Wing pouch regions of the discs expressing CD8-Cherry alone (top) or with AP-1μ RNAi (bottom) in the posterior compartment, visualised with E-cad-GFP (green, left), CD8 (magenta, left) and cleaved effector caspase (Dcp-1, yellow, left; inverted grayscale, right and bottom sagittal projection of the dashed line). Scale bars: 50 μm (top) and 20 μm (bottom). For AP-1 γ see Fig EV1H. **I** Basal region of wing discs expressing CD8-Cherry alone (top) or with AP-1μ RNAi (bottom) in the posterior compartment, co-stained with αPS1 integrin (green, left; inverted grayscale, right). Right panels display areas in the Anterior (A) and Posterior (P) compartments, which are highlighted by rectangles. Scale bars: 50 μm (left); 10 μm (right).

We observed complex morphological changes upon AP-1μ or AP-1 γ knockdown already in third instar wing imaginal discs (Fig 1B-E). Posterior compartments were reduced in size, as reflected by a smaller Posterior/Anterior (P/A) ratio than in controls (Fig 1B, D), while compartmental borders remained well defined (Fig 1E). Additionally, AP-1 knockdown resulted in ectopic folding of the wing pouch region (Fig 1E). While this folding could contribute to the reduced compartment size, we also examined for effects of AP-1 knockdown on cell size and number in the wing pouch. We found an increase in the apical area of the cells expressing the RNAis for the AP-1 subunits (Fig 1F-G). Combining this increase in the apical area with the reduction of total compartmental size indicates a reduced number of cells in the tissue following AP-1 knockdown.

The reduction in cell number could be a consequence of a reduced proliferation rate, increase in cell death, or extrusion of live cells. We found that the proliferation rate was however mildly increased when normalized to cell number, although the rate on the tissue level was unaffected (Fig EV1C-E). Concurrently, we detected a considerable amount of cell death visualised by the cleaved effector caspase Dcp-1 (Fig 1H, Fig EV1F). This increase in Dcp-1 signal was consistent with the presence of fragmented DAPI staining, indicative of nuclear debris (Fig EV1G). The basal localization of Dcp-1 and fragmented DAPI (Fig 1H and Fig EV1F, G) suggests that these cells are being eliminated from the tissue as described for dying cells (Bergantiños *et al*, 2010; Herrera *et al*, 2013). We found no evidence for extrusion of living cells.

Therefore, the three main effects of AP-1 knockdown in wing imaginal discs are: increased cell death, enlarged apical cell area, and ectopic folds. As the latter can be due to altered basal adhesion (Sui *et al*, 2018), we examined integrin localization upon AP-1 knockdown. Indeed, the αPS1 (Mew) integrin – specific for the dorsal compartment (Brower *et al*, 1984) – was nearly absent from the basal surface of the tissue (Fig 1I). We found similar changes in the Mew binding partner ßPS (Mys) and Laminin B2 (Fig EV2A), further supporting the loss of basal adhesions, and thus, tension. Mimicking the observed loss of basal adhesions by overexpressing diß, a chimeric protein acting as a dominant negative version of Mys (Martin-Bermudo *et al*, 1999; Domínguez-Giménez *et al*, 2007) (Fig EV2B-D), led to ectopic folds similar to AP-1 knockdown (Fig EV2C) (Domínguez-Giménez *et al*, 2007). In contrast, diß expression did not increase the apical cell size and caused only small pockets of dying cells (Fig EV2E-G). We concluded that downregulation of the AP-1 complex results in ectopic folds in wing pouch epithelium by interfering with basal localization of integrin complexes. This is likely due to incorrect trafficking of integrins from TGN leading to defective cell-to-extracellular matrix adhesions, altering the basal tension in the tissue. However, our findings suggest that AP-1 is involved in both determining apical cell area and promoting cell survival via a mechanism which is independent of integrins.

To gain insights into this mechanism, we next explored the intracellular localization of the AP-1 complex using the Venus-tagged AP-1μ subunit (AP-1μ-VFP) (Benhra *et al*, 2011). Consistent with observations in other cell types (Benhra *et al*, 2011; Burgess *et al*, 2011; Grant & Donaldson, 2009; Tan *et al*, 2019), AP-1μ-VFP accumulated within discrete puncta (spots) matching the TGN (Golgin-245-positive, (Riedel *et al*, 2016; Kondylis & Rabouille, 2009), Fig 2A) and REs (Rab11-positive, (Tanaka *et al*, 2008), Fig 2A). Unexpectedly, we also found a distinct subapical pool of AP-1μ colocalizing with E-cad at the AJs Fig 2A, B). We observed similar subapical pools of AP-1μ-VFP in embryonic epidermis and pupal eyes (Fig EV3A, B). In contrast, we detected only a small number of puncta at the basal surface, with no apparent co-localization with αPS1 (Fig 2A and Fig EV3C, (Brower *et al*, 1984)). Knockdown of the γ subunit of AP-1 reduced levels of AP-1μ-VFP both in the cytoplasm and at AJs (Fig 2C-E), which we measured as either the mean intensity of native VFP fluorescence (Fig 2D) or total content (Fig 2E) at either AJs or in the cytoplasm to account for the increased apical perimeter. These results suggested that the AP-1μ-VFP molecules assemble into the AP-1 complex at AJs as well as in the cytoplasm. This localization of AP-1μ-VFP at AJs also suggested that the complex might have roles directly there, consistent with the recent discovery of the mammalian AP-1B at the transient focal adhesions (Kell *et al*, 2020). We next sought to discover such roles by assessing the effects of AP-1 knockdown on endocytosis at the wing disc AJs through measuring the internalization of an antibody against the E-cad extracellular domain (Fig 2F, (Bulgakova & Brown, 2016)). To our surprise, we found that at 30 minutes after labelling the number of vesicles containing E-cad antibody almost doubled following AP-1μ knockdown in comparison to control (Fig 2G). The number of vesicles remained doubled at 60 minutes after labelling (Fig 2G). We conclude that E-cad internalization from AJs is elevated following AP-1 knockdown, indicating that the AP-1 complex negatively regulates E-cad endocytosis.

**Figure 2.**
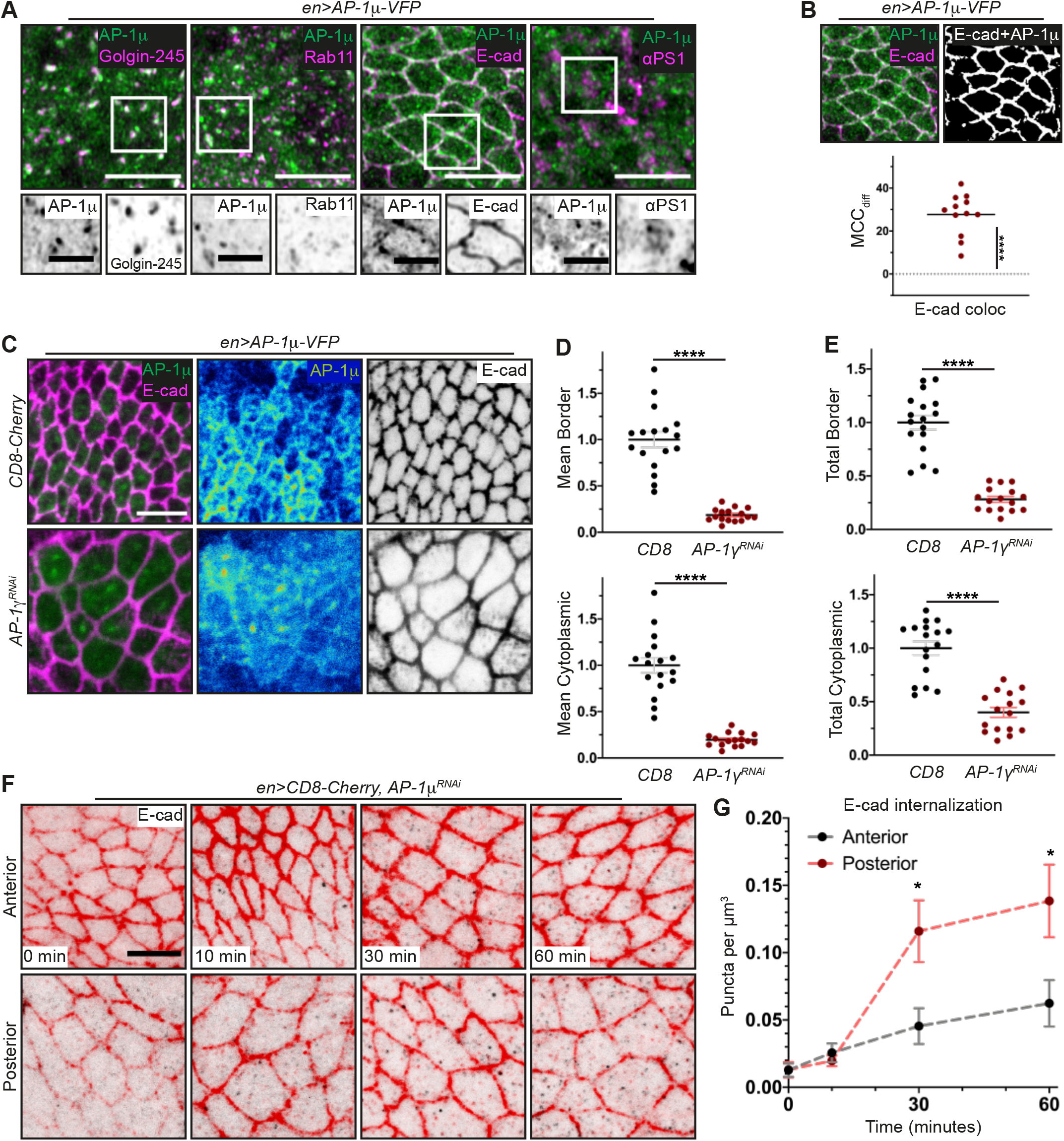
An apical fraction of AP-1 co-localises with E-cadherin and regulates its endocytosis. **A** Posterior compartments of wing discs expressing AP-1μ-VFP (green) and co-stained with intracellular markers (purple) of the Trans Golgi Network (Golgin-245), Recycling Endosomes (Rab11), the AJs (E-cad) and the basal membrane (integrin αPS1). Grayscale images correspond to the signal from AP-1μ-VFP and each marker within the white square depicted on the image above. Scale bars: 5 μm (top) or 2 μm (detail). **B** Example of colocalization (top) and the Manders’ Correlation Coefficient difference (MCC_diff_, see Methods) for the E-cad and AP-1μ-VFP threshold signal (bottom, n=12). ****P<0.0001 (one sample t-test in comparison to zero for random colocalization). **C** Posterior compartments of wing discs expressing AP-1μ-VFP (green, left; heatmap, right) together with CD8-Cherry (not shown, top) or AP-1γ RNAi (bottom) co-stained for E-cad to identify the AJs (magenta, left; and inverted grayscale, right). Scale bar: 5 μm. **D, E** Mean (D) and total (E) AP-1μ-VFP protein levels at the cell borders (top) and in the cytoplasm (bottom). Each dot represents cell average for an individual disc (n=17, 16). ****P<0.0001 (two-tailed t-test). **F** Pulse-chase labelling with E-cad antibody in the anterior (control, top) and posterior (expressing AP-1μ RNAi, bottom) compartments of wing discs. Apical region with AJs is shown in red, whereas the puncta in the 1.9 μm below (see Methods) are shown in black. Scale bar: 5 μm. **G** The number of vesicles per μm^3^ (mean ± s.e.m.) in the compartments and time points depicted in F (n=5-7 compartments/ time point). *P<0.05 (Welch’s t-test).

We next explored the consequences of the elevated internalisation of E-cad on its distribution at the AJs and intracellularly using the GFP-tagged E-cad driven by the ubiquitous *Ubi-p63E* promoter (*ubi*::E-cad-GFP)(Oda & Tsukita, 2001). We observed that the downregulation of both μ and γ subunits of AP-1 halved the levels of E-cad at the cellcell borders, which we measured as either the mean intensity of native GFP fluorescence at AJs or the total junctional content (Fig 3A,B, (Wu & Pollard, 2005; Coffman & Wu, 2012)). At the same time, while the mean signal of E-cad in the cytoplasm was also reduced, due to the increased apical cell area the total intracellular content remained normal (Fig 3C). We sought to verify this result using the knock-in of GFP into *shg* gene, producing E-cad-GFP protein expressed from the endogenous promoter (*shg*::E-cad-GFP, Fig 3D, (Huang *et al*, 2009)). Unexpectedly, the mean levels of *shg*::E-cad-GFP at AJs were only mildly reduced, while the total junctional protein content remained either unchanged or even elevated following AP-1 knockdown (Fig 3D-E). At the same time, the mean levels of *shg::* E-cad-GFP in cytoplasm were also only mildly reduced, with an increase of total intracellular protein content following knockdown of either AP-1μ or AP-1γ (Fig 3D,F). Therefore, for both E-cad-GFP proteins, knockdown of AP-1 elevated ratios of E-cad protein content in the cytoplasm relative to cell surface (Fig 3H, I), consistent with the increase in E-cad endocytosis (Fig 2F,G). We tested if this increase associated with E-cad retention at the REs/TGN, where the main pool of AP-1 localizes (Fig 2A). Using respective markers, we indeed found that AP-1μ knockdown increased co-localization of *ubi*::E-cad-GFP and *shg*:: E-cad-GFP with both organelles (Fig EV4A-F). These data suggest that AP-1 depletion increases the residence time of E-cad at both REs and TGN, whereas the negligible colocalization of E-cad with these organelles in control indicates that this residence time normally is very short.

**Figure 3.**
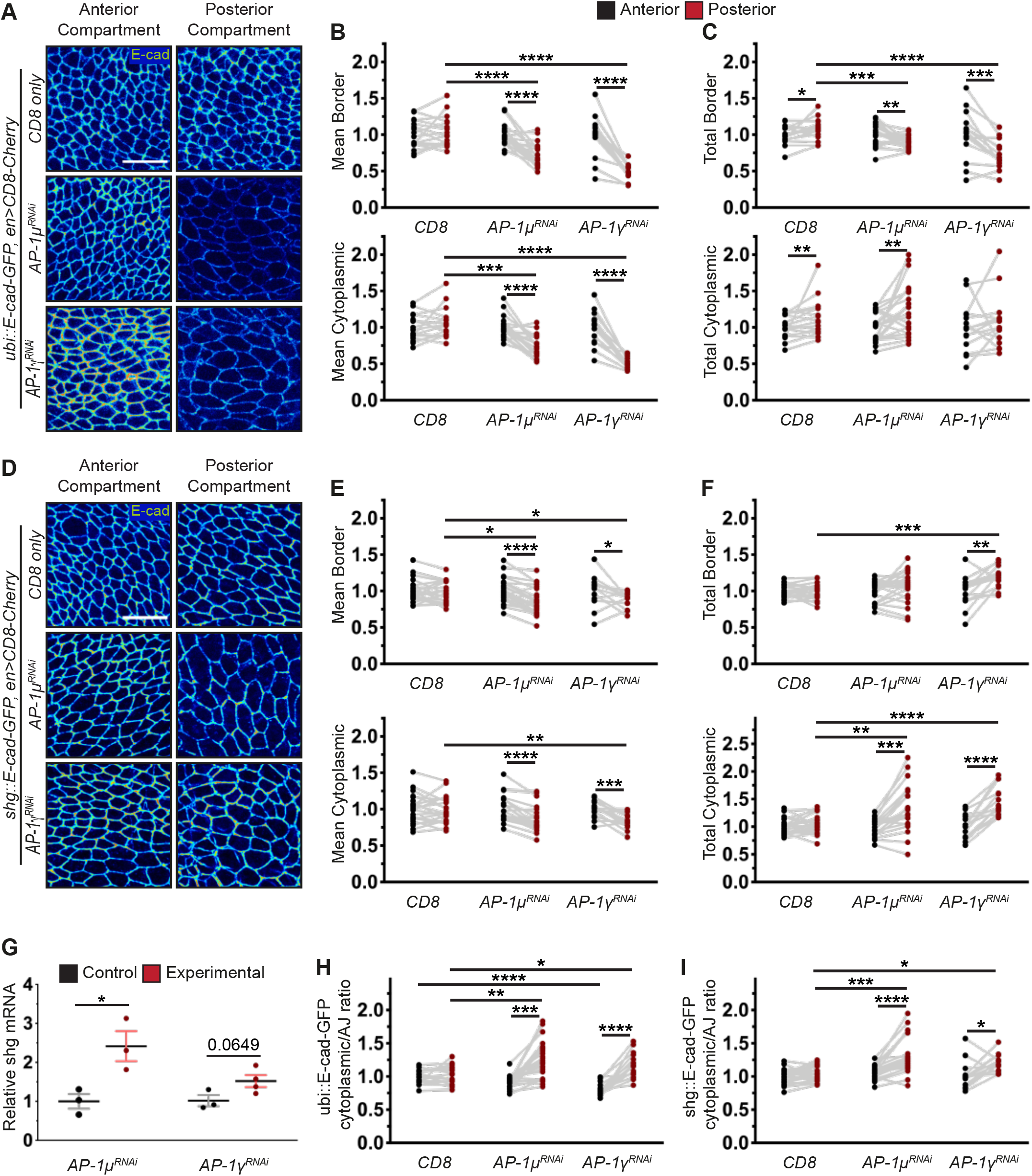
AP-1 controls the levels and the expression of E-cadherin. **A** Examples of cells in dorsal wing disc pouch expressing *ubi*::E-cad-GFP (heatmap) corresponding to anterior (left) and posterior (right) compartments of the indicated genotypes. Scale bar: 10 μm. **B, C** Mean (B) and total (C) *ubi*::E-cad-GFP protein levels at the cell borders (top) and in the cytoplasm (bottom). Each dot represents cell average for individual discs and compartments; compartments from same disc are connected by grey lines (n=17, 23, 15). **D** Examples of cells in dorsal wing disc pouch expressing *shg*::E-cad-GFP (heatmap) corresponding to anterior (left) and posterior (right) compartments of the indicated genotypes. Scale bar: 10 μm. **E, F** Mean (E) and total (F) *shg*::E-cad-GFP protein levels at the cell borders (top) and in the cytoplasm (bottom). Each dot represents cell average for individual discs and compartments; compartments from same disc are connected by grey lines (n=22, 23, 15). **G** Relative *shg* expression levels in control tissues and those with AP-1 subunits knockdown. Each dot represents an independent biological replicate (n= 3-4 per genotype). **H, I** Cytoplasmic/AJs ratio of the total protein levels for *ubi*::E-cad-GFP (H) and *shg*::E-cad-GFP (I). Datasets are same as above. Dots represent individual discs; compartments from same disc are connected by grey lines (n= 17, 23, 15 in H; 22, 23, 15 in I). *P<0.05, **P<0.01, ***P<0.001 and ****P<0.0001. Wilcoxon and paired t-test (anterior versus posterior) and Kluskal-Walls or two-Way ANOVA tests (comparisons between genotypes) were used in B, C, E, and F; Welch’s t-test – in G.

The two GFP-tagged E-cad variants differ in the promoters they are expressed from: a heterologous ubiquitous promoter or endogenous one. Therefore, the striking difference between the effects of AP-1 knockdown on resulting protein amounts could be explained at the promoter level – namely that AP-1 knockdown increases *shg* gene expression. We tested this possibility using RT-qPCR and found that the downregulation of the AP-1 complex increased *shg* mRNA levels (Fig 3G). This suggested the existence of a yet-to-characterise feedback loop whereby cells upregulate expression of E-cad in response to a perturbation in trafficking following AP-1 knockdown. Altogether, we suggest that AP-1 knockdown results in both enhanced internalization of E-cad from the cell surface and retention of this internalized E-cad at TGN and REs, leading to the activation of the feedback mechanism (Fig 4A). Such a mechanism has not been reported in the past to our knowledge and provides a novel layer of cell-cell adhesion regulation. We speculate that this mechanism ensures tissue resilience to stochastic defects in E-cad trafficking.

**Figure 4.**
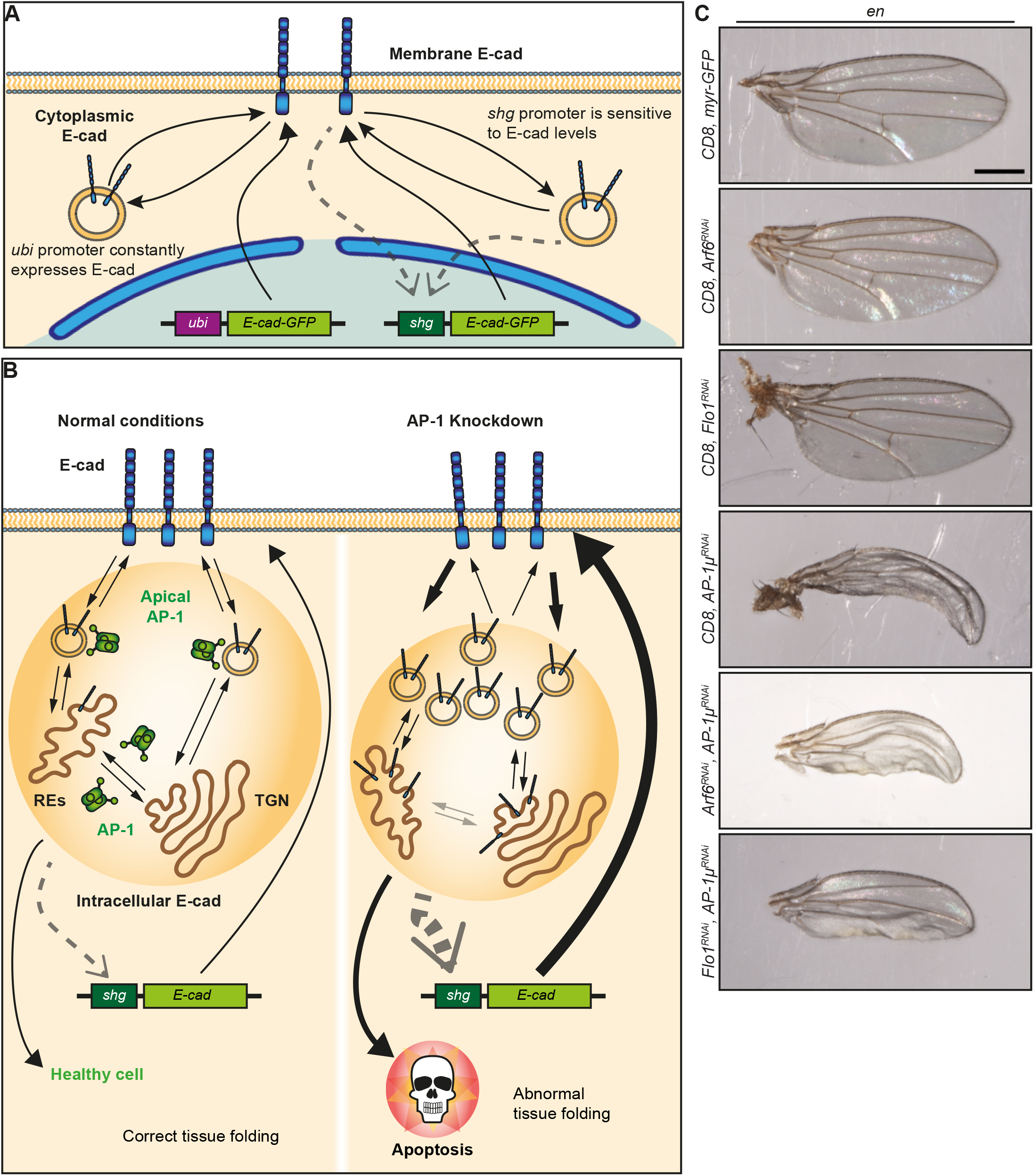
AP-1 promotes cell survival by controlling intracellular levels of E-cadherin. **A** Model of the proposed E-cad feedback loop. The ratio between the membrane and intracellular levels of E-cad influences the transcription of the *shg* gene. This leads to different outcomes for the heterologously expressed *ubi*::E-cad-GFP and the endogenous *shg*::E-cad-GFP, as only the latter is affected by this change in expression. **B** Model of the functions of AP-1 in the developing wing discs. In normal conditions (left), besides vesicle trafficking between the Recycling Endosomes (REs) and the Trans Golgi Network (TGN), an apical fraction of AP-1 remains at/near the E-cad-positive AJs, where it limits internalisation of E-cad. Upon AP-1 knockdown (right), E-cad endocytosis is elevated while E-cad is simultaneously retained in TGN/REs. This increases E-cad intracellular levels and induces cell death. At the same time, to bring E-cad membrane levels back to normal, cells increase *shg* expression. **C** Adult female fly wings expressing the indicated proteins or RNAis in their posterior compartments. Scale bar: 0.5 mm.

While the exact effects of AP-1 knockdown on levels of *shg::* E-cad-GFP and *ubi*::E-cad-GFP were different, both proteins displayed increased cytoplasmic levels and elevated cytoplasm/AJs ratio (Fig 3H, I). Therefore, we asked if the elevated E-cad internalization and intracellular retention were responsible for apoptosis following AP-1 knockdown (Fig 1H and 4B), as in the wing disc the failure to localize E-cad at AJs induces apoptosis through activation of JNK signalling (Jezowska *et al*, 2011). We recently showed that Flotillin1 promotes E-cad endocytosis via recruitment of the small GTPase Arf6 to AJs (Greig & Bulgakova, 2020). As Arf6 interacts with and regulates AP-1 (Tan *et al*, 2019; Shteyn *et al*, 2011), we tested whether Flotillin1 and Arf6 might act upstream of AP-1 at AJs. To this end, we downregulated either Arf6 or Flotillin1 simultaneously with AP-1, balancing the UAS-Gal4 dosage in controls with CD8-Cherry (Fig 4C). Indeed, both Flo1 or Arf6 knockdowns ameliorated the effects of AP-1 knockdown – adult wings had detectable posterior compartments with wing blisters (Fig 4C). These data demonstrate that inhibition of E-cad endocytosis rescues tissue viability but not the blister-producing loss of integrin adhesion (Domínguez-Giménez *et al*, 2007; Bökel & Brown, 2002). Furthermore, these results support that it is the elevated cytoplasmic E-cad which is inducing cell death (Fig 4B). We speculate that such a mechanism would be tumour-suppressive, as it will eliminate cells which hyperinternalize E-cad to undergo unregulated epithelial-to-mesenchymal transition.

In summary, we demonstrate that a newly discovered subapical pool of the AP-1 complex exerts a brake upon E-cad internalization, excess of which promotes cell death. We provide evidence that this AP-1 function is accompanied by a transcriptional feedback which maintains E-cad levels at the cell surface. Finally, our data support the role of the small GTPase Arf6 for the AP-1 function at the plasma membrane. In line with the correlation of the loss of tissue-specific AP-1B with epithelial architecture and the metastatic potential in humans (Kell *et al*, 2020), our findings highlight the pivotal role of the AP-1 complex in preventing tumour progression but also enabling correct epithelial morphogenesis.

## Materials and Methods

### Reagents and tool table

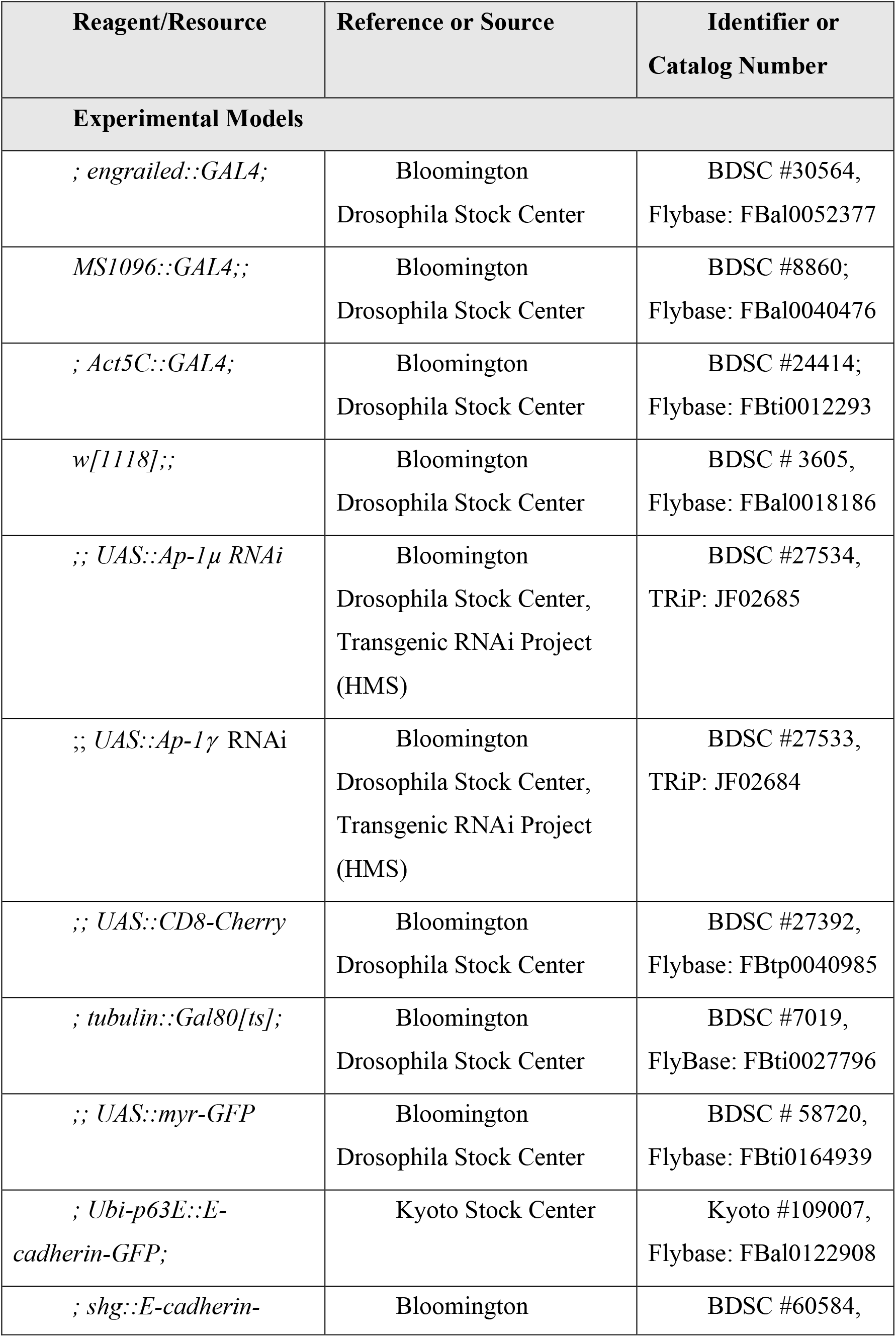
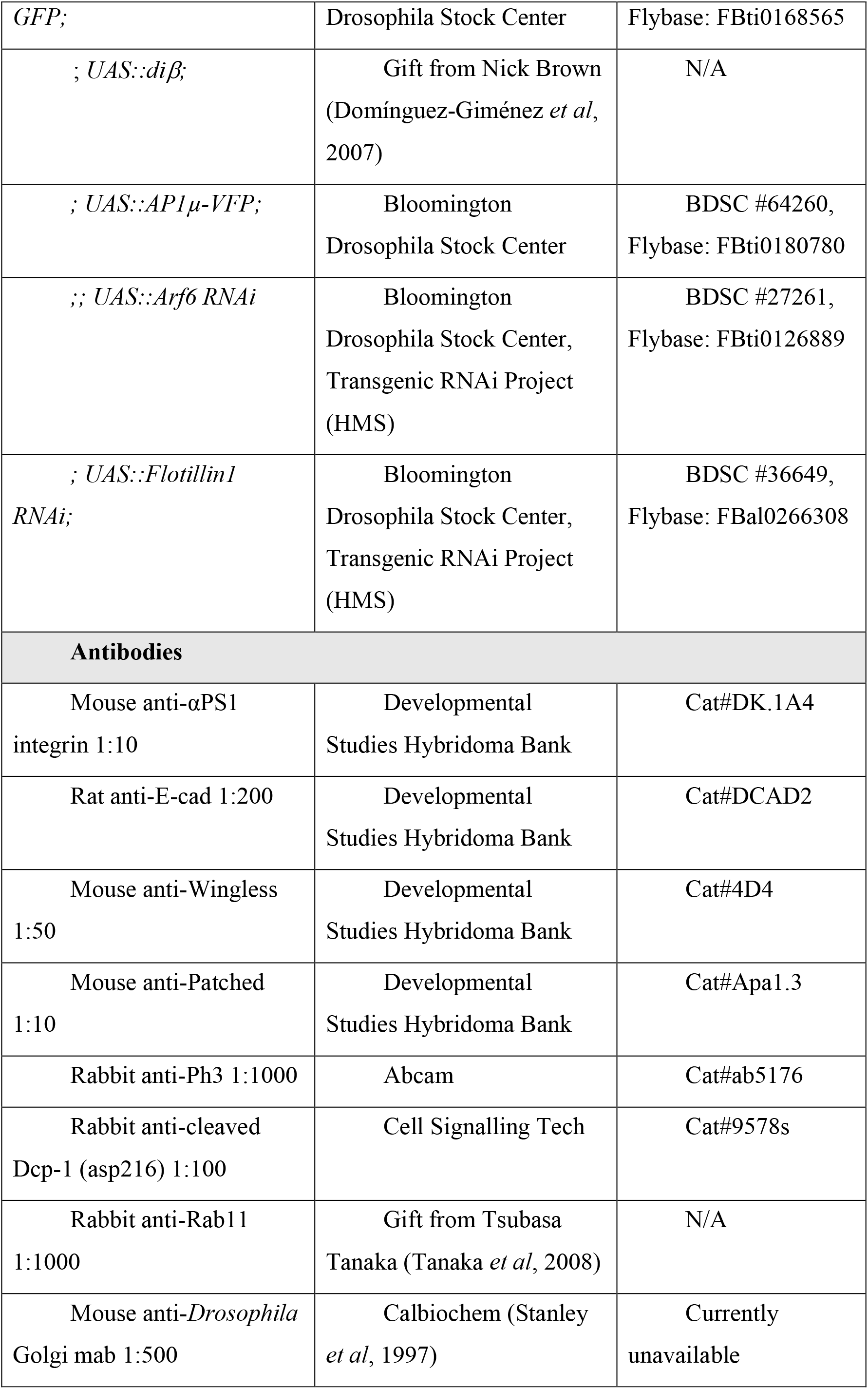
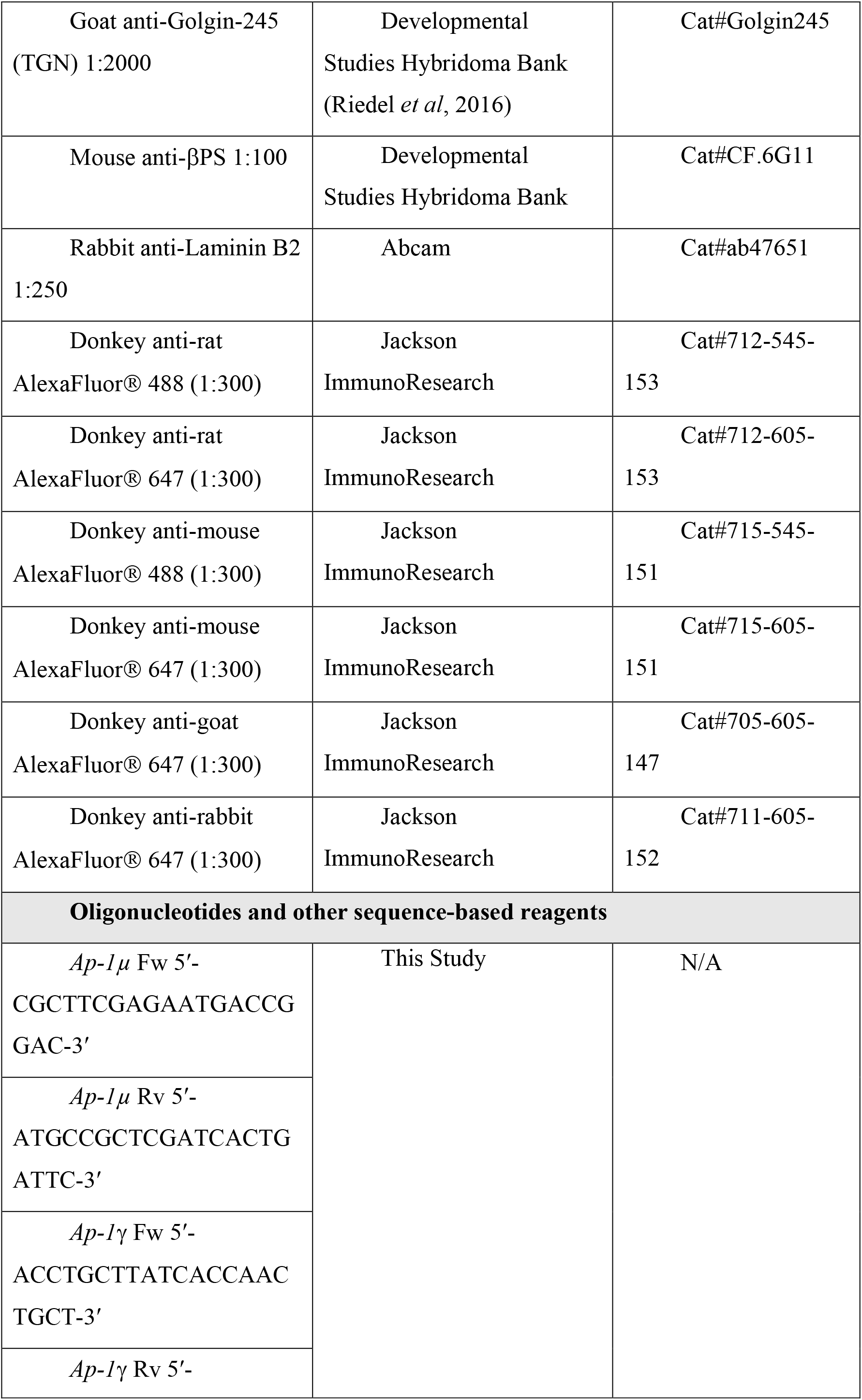
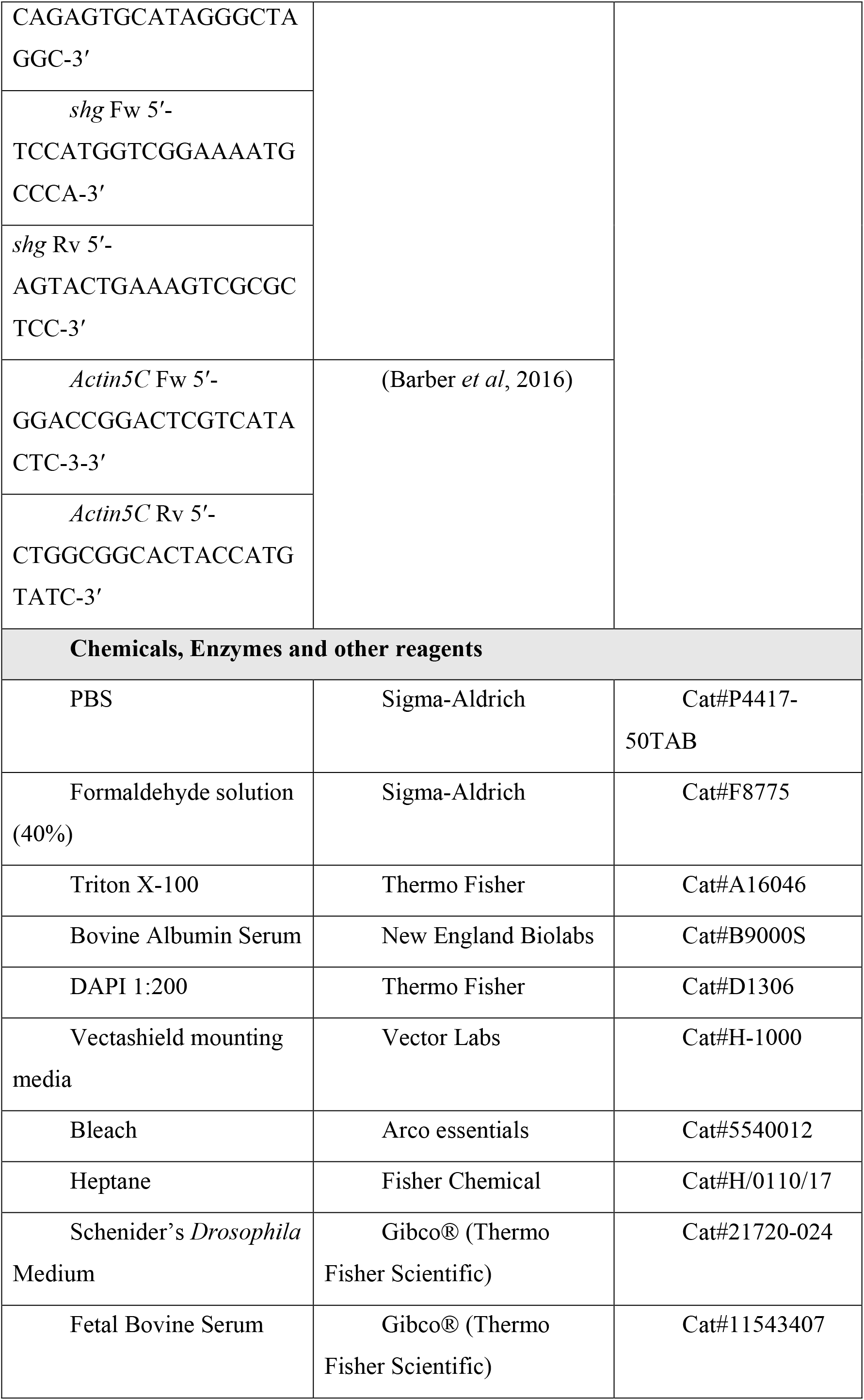
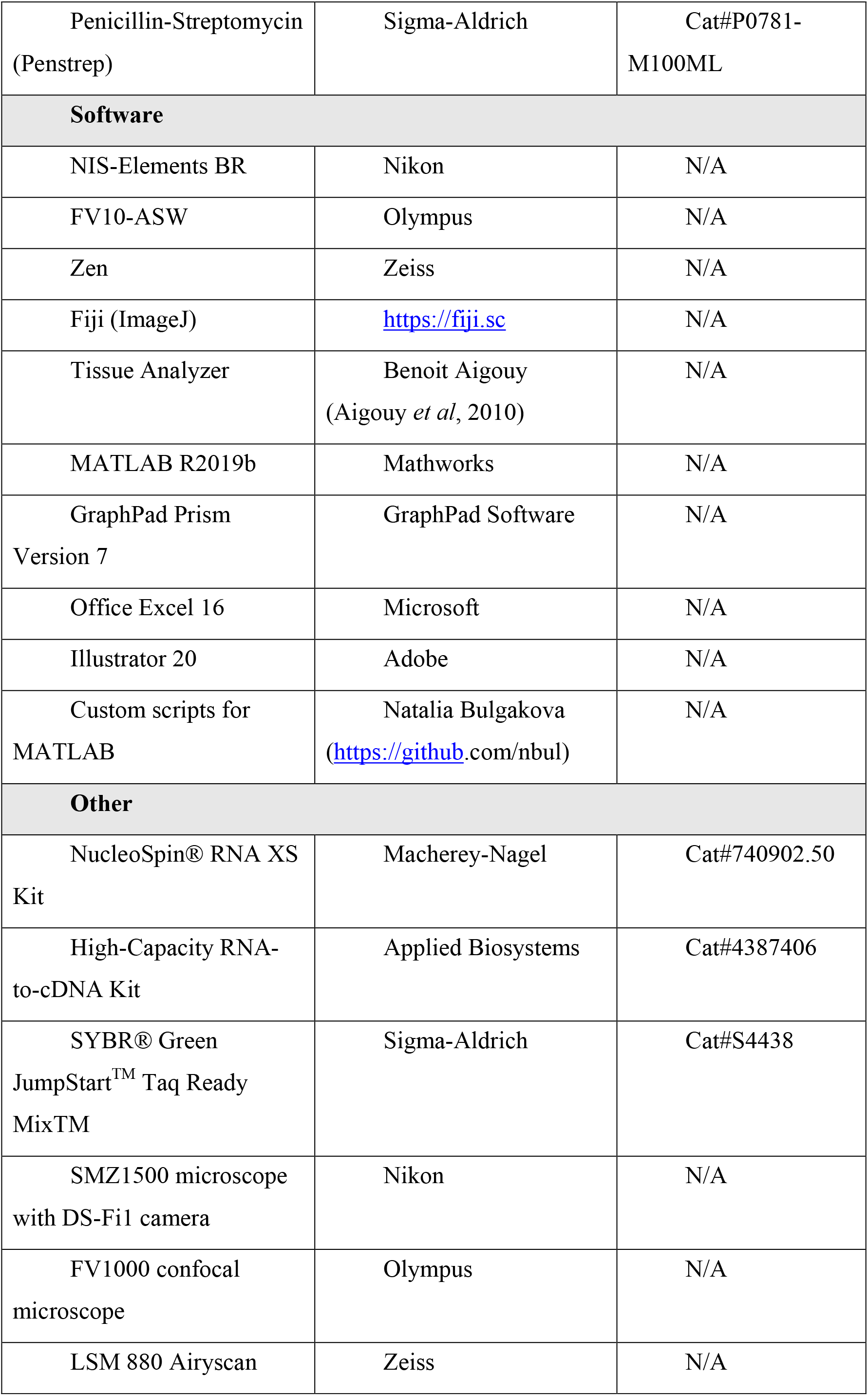
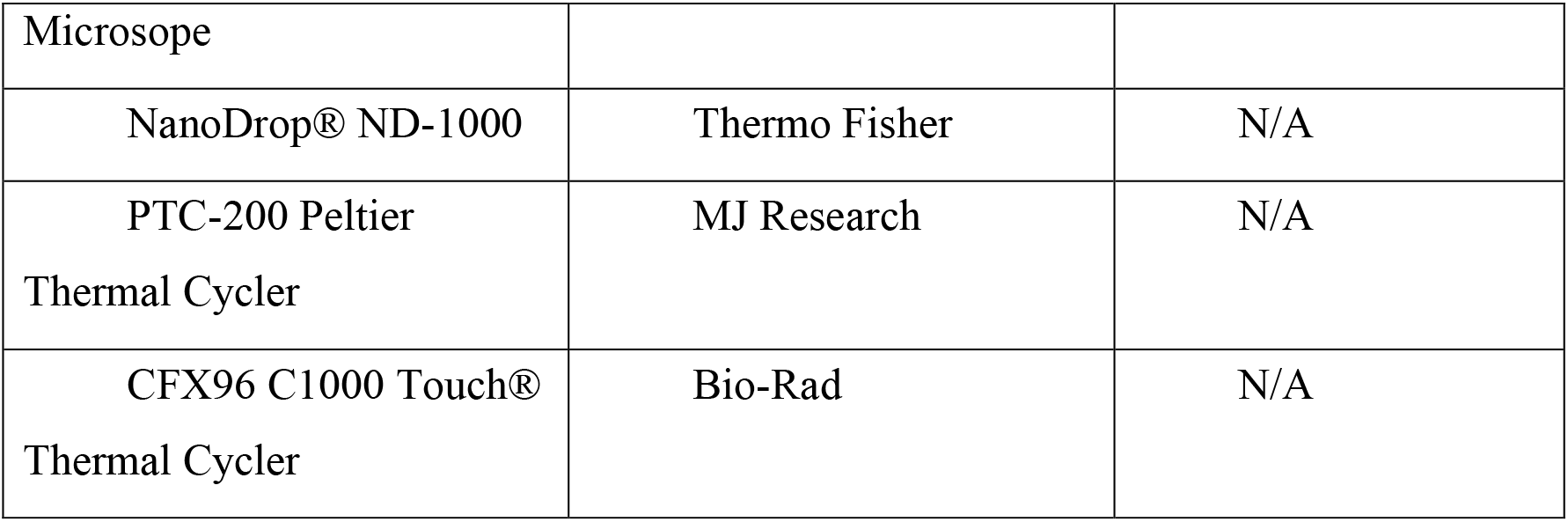

## Methods and Protocols

### Fly genetics and husbandry

*Drosophila melanogaster* were raised on standard cornmeal/agar/molasses media at 18°C or 25°C unless otherwise specified. To express constructs of interest, the GAL4/UAS system was used (Brand & Perrimon, 1993), with *engrailed::GAL4* (*en*::*Gal4*), *MS1096::GAL4*, or *Act5C::GAL4* drivers. To examine adult wings, flies were raised at 18°C to circumvent potential lethality due to RNAi expression. To study larval wing discs, larvae were raised at 25°C. The GAL4:UAS ratio was kept constant within each experimental dataset using additional copies of *UAS*::CD8-Cherry or *UAS*::myristoylated-GFP (*UAS*::myr-GFP). Acute expression of RNAi was achieved with the combination of *Act5C*::GAL4 and *twbwlin*::GAL80^ts^. Larvae then were raised at 18°C for thirteen days after egg laying and shifted to 29°C for 48 hours prior to dissection of the wing discs.

### Adult wings imaging

Female adult flies were frozen upon collection with CO2, and wings were removed and imaged with a Nikon SMZ1500 microscope equipped with Nikon DS-Fi1 camera controlled by NIS-Elements BR software.

### Dissection and Immunostaining

#### Wing discs

Third instar larvae were dissected after being kept for six days after egg laying at 25°C. Cuticles with attached imaginal discs were fixed for 15 minutes with 4% formaldehyde (Sigma, F8775) in PBS (Phosphate Buffer Saline, Sigma-Aldrich, P4417) at room temperature, then washed with PBS with 0.1% Triton X-100 (Thermo Fisher, A16046, hereafter PBST), and incubated with 1% Bovine Serum Albumin (BSA, New England Biolabs, B9000S) in PBST for 1 hour at room temperature. Cuticles were incubated with primary antibodies and 1% BSA in PBST overnight at 4°C, washed in PBST, incubated overnight at 4°C with secondary antibodies and 1% BSA in PBST (including DAPI when applicable), and washed in PBST again. All antibodies and their concentrations are listed in the Reagents and Tools Table. Finally, discs were separated from the cuticles and mounted in Vectashield (Vector Labs, H-1000). For samples not requiring immunostaining, e.g. imaging direct fluorescence of E-cad-GFP, discs were immediately mounted following fixation and one round of washing in PBST.

#### Embryos and retinas (Fig EV2)

*Drosophila* embryos were aged up to the desired developmental stage with 3-hour collections at 25°C, followed by 21 hours at 18°C. Embryos were then dechorionated in 50% commercial bleach solution in water for four minutes. Following extensive washing with deionised water, embryos were fixed with a 1:1 4% formaldehyde in PBS: heptane (Fisher Chemical, H-0110-17) for 20 minutes with constant agitation at room temperature. Embryos were devitellinized by vigorous shaking in 1:1 methanol: heptane for 20 seconds, washed and stored in methanol at −20°C upon required. Methanol was removed by washing in PBST, and embryos were subjected to the same staining procedure as the wing discs, but using 0.05% Triton in PBS rather than 0.1%.

Prepupal stage individuals raised at 25°C were collected and aged for 40 hours at 25°C to reach the desired developmental stage. Retinas were extracted by cutting the external cuticle open in PBS. The same protocol as for wing discs was followed for staining.

#### Pulse-chase assay

Wing discs were dissected in Schneider’s Insect medium (Gibco, 21720-024) with 1% Penstrep (Sigma-Aldrich, P0781-M100ML) and 5% Fetal Bovine Serum (FBS, Gibco, 11543407), and the peripodial membrane was removed with a fine needle. Then, the discs were incubated with rat anti-E-cad antibody (DCAD2, 1:200, DSHB) in the same medium at 4°C for 1 hour. After three quick washes with cold medium, the discs were incubated in fresh medium at room temperature. Several (5-7) discs were fixed in 4% formaldehyde in PBS for 15 minutes at each analysed time point (immediately after washing for t = 0 min). The discs were stained as described above using AlexaFluor^®^ 488-conjugated anti-rat antibody (Jackson ImmunoResearch).

### Microscopy, data acquisition and image processing

All microscopy experiments except colocalization analyses were done using an upright Olympus FV1000 confocal microscope with either 60x/1.40 NA (fluorescence intensity and cell morphology) or 20x/0.75 NA (tissue morphology) objectives. In the former case, images were taken in the dorsal region of the wing disc pouch (Fig 1B): for each disc and compartment a z-stack of 6 slices spaced by 0.38 μm was acquired capturing the complete span of the apical Adherens Junctions. This z-spacing was used for all the acquisitions with the 60x objective, while 0.80 μm spacing was used for z-stacks done with the 20x objective. All the images were at 16-bit depth in Olympus binary image format.

Images used for colocalization assays were taken using an inverted Zeiss LSM 880 Airyscan confocal microscope. Z-stacks of 20 slices spaced by 0.38 μm (AP-1mu-VFP colocalization) or 10 slices spaced by 0.18 μm (E-cad co-localization) were taken with a 63x/1.40 NA objective. Raw 16-bit images were processed using Zeiss software (automatic mode) to obtain “.czi” files.

#### Compartment size

Z-stacks with E-cad-GFP and CD8-Cherry signals were projected using the “maximum intensity” algorithm in Fiji (https://fiji.sc) and used to measure the area of the whole disc and posterior compartment, respectively, with the Fiji selection tool. The anterior compartment area was calculated by subtracting the posterior compartment area from that of the disc.

#### Apical cell area, membrane and cytoplasmic protein levels

For the analyses of membrane and cytoplasmic protein levels, z-stacks with signal of GFP fluorescence were projected using the “average intensity” algorithm in Fiji. To distinguish membrane and cytoplasm as well as measure apical cell area, we generated binary masks from images with visualized cell outlines (i.e. E-cad signal). In particular, z-stacks with E-cad signal were projected using the “maximum intensity” algorithm in Fiji. The background was subtracted from these projections using a rolling ball of 50-pixel radius, and their brightness and contrast were adjusted using the automatic optimization algorithm in Fiji. These projections were used to generate masks using the Tissue Analyzer plugin in Fiji (Aigouy *et al*, 2010). Cells in which the AJs were not completely in focus were removed manually.

The binary masks were used to measure the fluorescence intensity of average projections using our in-house Matlab script (https://github.com/nbul/Intensity). First, individual objects (cells) and their boundaries were identified from each binary mask. Then the identified objects (cells) were analysed on a cell-by-cell basis. The area of objects and length of their boundaries in pixels were determined, and then manually converted from pixels to μm^2^. The boundary was dilated using a diamond-shaped morphological structural element of size 3 to encompass the XY spread of E-cad signal. The mean and total (sum of all pixel intensities) intensities of the dilated boundary (membrane signal) and the object with subtracted boundary (cytoplasm) were calculated. All the values were averaged to produce single values per wing disc thus testing biological replicates and excluding chances of one disc having higher contribution to the result due to variable number of cells in each disc.

#### Proliferation

z-stacks with E-cad-GFP and pH3 (647) signal corresponding to the anterior and posterior dorsal wing pouches were processed separately for each channel. E-cad signal was segmented as described above using Tissue Analyzer. The resulting binary masks were dilated and then inverted. The area with cells was used then to generate maximum projections of the equivalent region for the pH3 channel. The projection was processed using the following steps: background was subtracted using a rolling ball of 1-pixel radius; then Gaussian blur of 2-pixel radius was added; and image was binarized using a threshold set to three times the average intensity of the original maximum projection.

To calculate the cell number and the total cell area in the imaged regions from the processed binary segmented masks, we employed the Particle Analysis plugin on Fiji, limiting particle-detection to sizes up to 67.5 um^2^ (or 3000 pixel^2^). The number of proliferating cells in the same area was determined using the processed images of the pH3 channel corresponding to the same area using particle sizes between 5 and 50 um^2^. These numbers were used to calculate the number of dividing cells per 100 cells or per 100 um^2^.

#### Co-localization

For co-localization of AP-1μ-VFP with cellular markers, we employed the co-localization plugin Coloc 2 in Fiji (https://imagej.net/Coloc2). We selected Manders’s Colocalization Coefficients (MCC (Manders *et al*, 1993; McDonald & Dunn, 2013)) as more informative for probes distributed to more than one compartment than Pearson’s correlation coefficients (Dunn *et al*, 2011). The MCCdiff value was obtained by subtracting the percentage of pixels positive for AP-1μ-VFP (obtained using the threshold determined by Coloc 2) from the percentage of pixels positive for the marker that were also positive for AP-1μ-VFP.

For co-localization of E-cad-GFP with markers of intracellular organelles, we analysed the images with an in-house script at MATLAB (https://github.com/nbul/Localization). This script followed the same principle to obtain MCCdiff values as the Coloc 2 plugin, but is more versatile as it enables a manual selection of a threshold method for each probe. Such selection is required as the bright signal of E-cad-GFP at cell borders limits its detection in cytoplasm using standard threshold methods. The E-cad-GFP signal at cell borders was excluded from the analysis, and the resulting MCCdiff values accounted only therefore for the intracellular signal.

#### Pulse-chase

12 sections, comprising 6 sections spanning the complete region of Adherens Junctions visualized with the staining of the antibody bound to E-cad at the cell surface and the 6 sections immediately basal to Adherens Junctions, were used for counting the E-cad-positive vesicles. Firstly, a binary mask with cell outlines was created from a projection of the 6 apical-most sections as described above. The average cytoplasmic signal and cell area were measured with our in-house script as described above and used to threshold the corresponding maximum projection of the 6 basal sections in Fiji using the following steps: background was subtracted using a rolling ball of 4-pixel radius; then Gaussian blur of 2-pixel radius was added; and image was binarized using a threshold set to 70% of the average cytoplasmic signal of the apical section. The resulting images were used for the analysis using the Particle Analysis plugin in Fiji, limiting particle-detection to sizes between 0.05 and 0.45 um^2^. Puncta density was determined using the average apical area of the cells.

#### Other processing

Sagittal views of the wing discs, for example in Fig 1, were generated using the “Reslice” tool in Fiji. For representative cases shown in Figures, maximum projections of the regions of interest were generated in Fiji using minimum modification, such as tilting, cropping, or automatic contrast of the whole view, for better presentation.

### RQ-qPCR

Primers for RT-qPCR were designed and *in silico* tested using Flybase (https://flybase.org), Primer-BLAST (https://www.ncbi.nlm.nih.gov/tools/primer-blast/index.cgi), and Net Primer (http://www.premierbiosoft.com/netprimer/) tools, and whenever possible aimed to target sequences separated by introns and present in all the splicing variants. All primers and amplicon sizes are listed in the Resources and Tools Table, and were manufactured by Thermo Fisher. RNA was extracted from wing discs dissected on ice using the NucleoSpin^®^ RNA XS Kit (Macherey-Nagel, 740902.50). Control wing discs expressed *UAS*::myr-GFP instead of the RNAi. For AP-1γ knockdown, only discs from male larvae were used to exclude the potential effects of dosage compensation. RNA concentration was determined with an ND-1000 Spectrophotometer (Nanodrop^®^, Thermo Fisher) and immediately used to generate cDNA with the High-Capacity RNA-to-cDNA Kit (Applied Biosystems, 4387406) using 400 ng of RNA. cDNA was then used as a template for the quantitative PCR reaction using SYBR^®^ Green JumpStart™ Taq Ready Mix™ (Sigma-Aldrich, S4438) using a CFX96™ Real-Time System (Bio-rad) and its proprietary software. All the primers were tested with standard reaction curves and gel electrophoresis. Upon testing of several widely employed housekeeping genes in *Drosophila* (Ponton *et al*, 2011; Lü *et al*, 2018), primers against *Actin5C* were selected due to their reproducible performance. Reactions were carried out in 3 technical replicates per a biological replicate (15 wing discs), in a volume of 10 μl with a primer concentration of 1 pmol/μl. At least three biological replicates were done per genotype. Ct values were obtained from SYBR fluorescence using thresholds determined from the standard curves (Larionov *et al*, 2005). Primer purity was tested on a control without any template in every performed assay.

Expression levels were determined by the 2^-ΔΔCT^ method (Schmittgen & Livak, 2008). For each biological replicate, Ct values were averaged across all technical replicates; and the average Ct value of *Actin5C* was subtracted from the target gene. This result (ΔCT) was normalized by subtracting the average ΔCT of the control genotype, producing ΔΔCT. This value was converted using the formula 2^-ΔΔCT^.

### Statistical Analysis

All the statistical analyses were done in GraphPad Prism 7 (https://graphpad.com/scientific-software/prism/). First, the datasets were cleaned from outliers using the ROUT detection method, and the distributions were tested for being normal with D’Agostino & Pearson test.

Precise n numbers, the type of statistical test, and the type of represented data (i.e. individual cases, Mean and SD, etc…) are described in the Figure legends. Significance was visually depicted in all the graphs with either the precise value or asterisks (* – P<0.05, ** – P<0.001, *** – P<0.0001, and **** – P<0.00001). For commonly used tests, such as t-test, two-tailed versions were used. Non-parametric tests were used when at least one sample did not display normal distribution, and appropriate corrections were applied if the assumption of equality of standard deviations was not met.

#### Posterior.anterior compartment size ratio

Posterior:anterior compartment size ratios of discs expressing RNAi against subunits of the AP-1 complex were tested against the external control discs expressing CD8-Cherry only using the Brown-Forsythe and Welch ANOVA test.

#### Proliferating cells

The number of dividing cells per 100 cells or per 100 um^2^ were compared with paired t-test/Wilcoxon test (between compartments of the same genotype) and Kruskal-Wallis test (between genotypes and the anterior control compartment).

#### Protein levels and cell size

Differences between genotypes were tested with two-way ANOVA (normal distribution) or Kluskal-Wallis test. Differences between the paired sets of anterior and posterior compartments were tested with paired t-test (normal distribution) or Wilcoxon test.

#### Co-localization

The differences between control and experimental compartments were tested using paired t-test (normal distribution) or Wilcoxon test. The MCCdiff value was also tested against zero (random colocalization (McDonald & Dunn, 2013)) using the one sample t-test.

#### Pulse-chase

The puncta density was compared between the compartments using Welch’s t-test.

#### Expression analyses

The gene expression in RNAi-expressing samples was compared to myr-GFP-expressing controls using Welch’s t-test.

## Data availability

### Materials Availability

This study generated new reagents by genetic recombination of available *Drosophila* strains, which are available upon request.

### Data and Code Availability

All in-house scripts for image analyses used in this study are available at https://github.com/nbul/

## Acknowledgements

We thank Nick Brown, Kyra Campbell, Iwan Evans, David Strutt, and Tsubasa Tanaka for fly stocks and antibodies; the University of Sheffield’s Wolfson Light Microscopy and Fly Facilities for their technical support; and Kyra Campbell and David Strutt for critical feedback on the manuscript. This work was supported by a grant from the UK Biotechnology and Biological Sciences Research Council (BB/P007503/1) to N.A.B.; a summer studentship from the Genetics Society and Think Ahead SURE scheme (University of Sheffield) to M.R.M. and K.B.

## Author Contributions

Conceptualization: N.A.B.; Methodology: M.R.M., N.A.B.; Software: N.A.B.; Validation: M.R.M.; Formal analysis: M.R.M., K.B., N.A.B.; Investigation: M.R.M., K.B.; Resources: M.R.M., N.A.B.; Data curation: M.R.M., N.A.B.; Writing – Original draft: M. R.M., N.A.B.; Writing – Review & editing: M.R.M., N.A.B.; Visualization: M.R.M., N. A.B.; Supervision: M.R.M., N.A.B.; Project Administration: N.A.B.; Funding acquisition: N.A.B.

## Expanded View Figure Legends

**Figure EV1.**
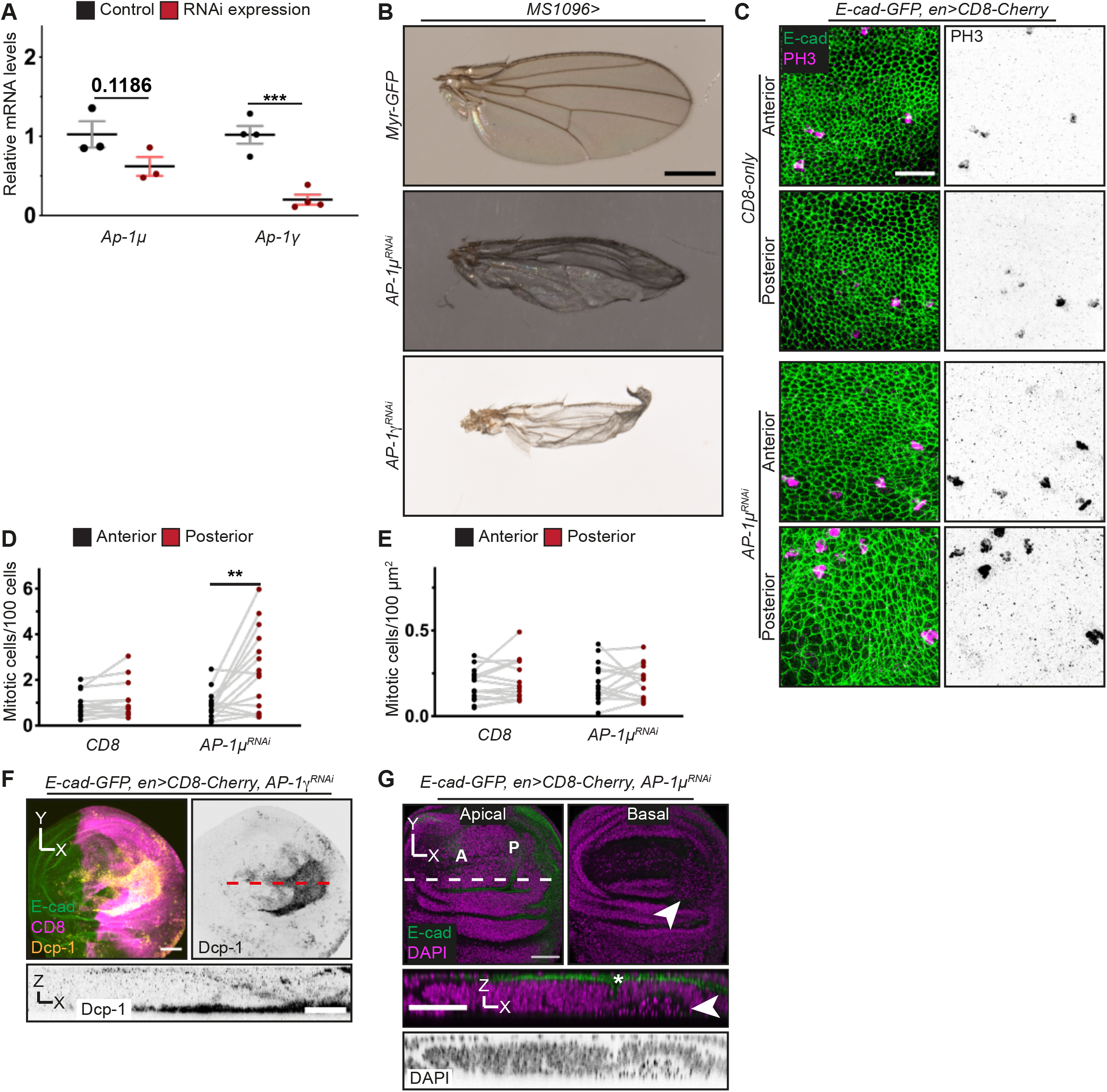
AP-1 regulates tissue development, cell morphology and cell survival at the wing disc (related to Figure 1). **A** Relative *AP-1μ* and *AP-1γ* expression levels in control tissues and those with RNAi knockdown. Each dot represents an independent biological replicate (n=3-4 per genotype). ***P<0.05 (Welch’s t-test). **B** Adult female fly wings expressing either myr-GFP or AP-1μ RNAi or AP-1γ RNAi in the wing pouch. Scale bar: 0.5 mm. **C** Examples of cells in dorsal anterior (left panels) and posterior (right panels) compartments of wing discs expressing either CD8-Cherry protein (not shown) alone (top) or with AP-1μ RNAi (bottom) in the posterior compartment, stained for E-cad (green, left) and the mitotic marker phospho-Histone H3 (PH3, magenta and inverted grayscale). Scale bar: 15 μm. **D, E** Proliferating cells per each 100 cells (D) and 100 μm^2^ of apical surface (G) in the dorsal wing disc pouch. Each dot represents the ratio for individual discs and compartments (n=16, 15). **P<0.01 (Wilcoxon test). **F** Wing pouch regions of a disc expressing CD8-Cherry (not shown) with AP-1μ RNAi in the posterior compartment, visualized with E-cad (green, top) and stained with DAPI (magenta and inverted grayscale in sagittal projection of the dashed line). Top panels display apical (left) and basal (right) areas in the Anterior (A) and Posterior (P) compartments. Arrowheads indicate basal accumulation of nuclear debris in the posterior compartment. Asterisk indicates the ectopic fold (related to Fig 1E). Scale bars: 50 μm (top) and 20 μm (bottom). **G** (related to Fig 1H) Wing pouch regions of a disc expressing CD8-Cherry with AP-1γ RNAi in the posterior compartment visualised with E-cad-GFP (green, left), CD8-Cherry (magenta, left) and cleaved caspase-3 (DCP-1, yellow, left; inverted grayscale, right and bottom sagittal projection of the dashed line. Scale bars: 50 μm (top) and 20 μm (bottom).

**Figure EV2.**
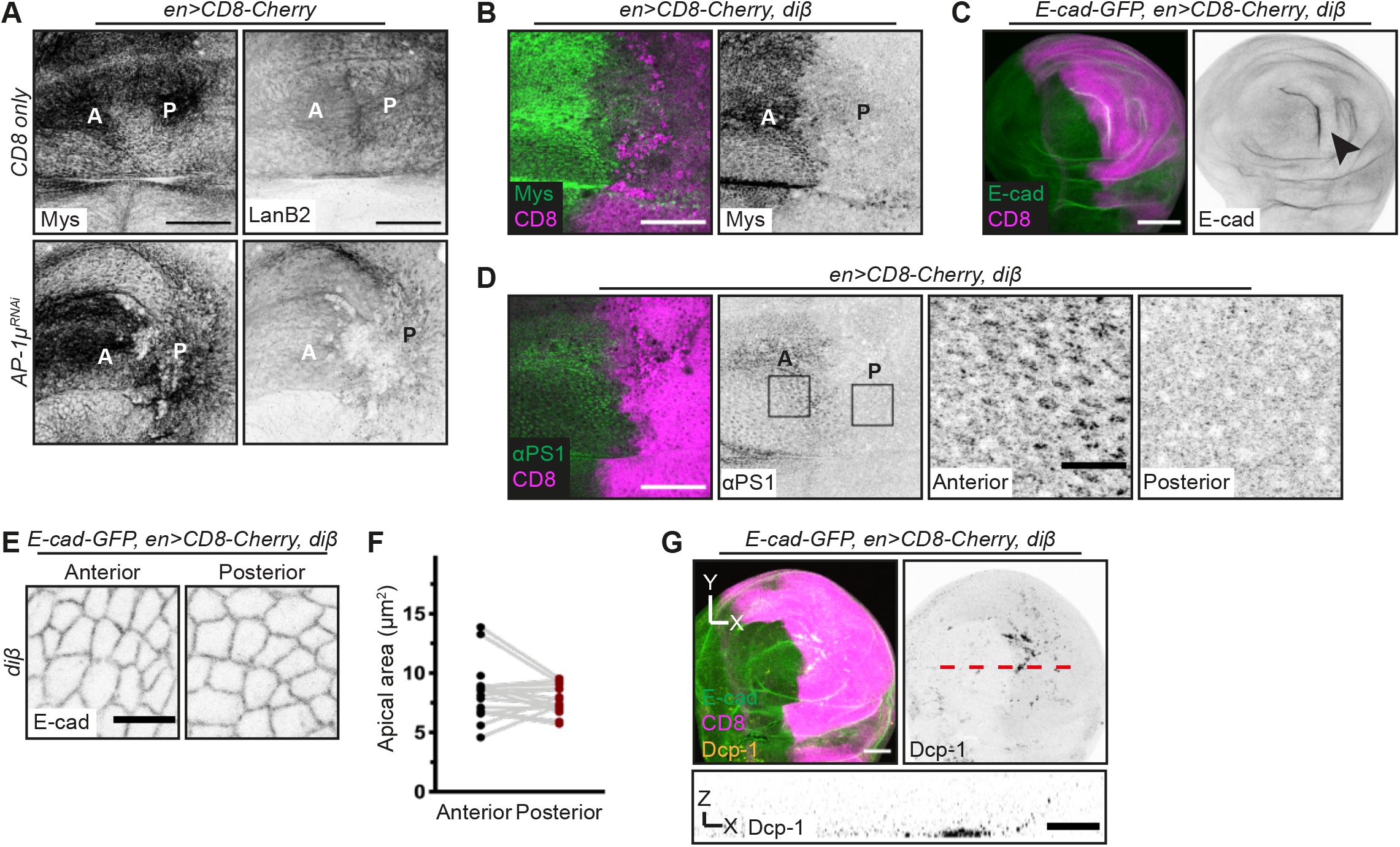
Disruption of basal polarity is not responsible for the AP-1 knockdown phenotype (related to Figure 1). **A** Basal region of wing discs expressing CD8-Cherry (not shown) alone (top) or with AP-1μ RNAi (bottom) in the posterior compartment (P, with anterior compartment, A, as internal control), stained for ßPS integrin (Mys, left) or Laminin B2 (LanB2, right). Scale bar: 50 μm. **B** Basal region of wing disc expressing CD8-Cherry (magenta, left) and diß protein at the posterior compartment (P, with anterior compartment, A, as internal control). The staining for ßPS integrin (Mys, green, left; inverted grayscale, right) confirms the displacement of the endogenous ßPS. Scale bar: 50 μm. **C** Wing pouch regions of discs expressing CD8-Cherry protein (magenta, left) with diß in the posterior compartment. E-cad-GFP is shown to visualize tissue architecture (green, left; inverted grayscale, right). Arrowhead indicates an ectopic fold. Scale bar: 100 μm. **D** Basal region of a wing disc expressing CD8-Cherry with diß in the posterior compartment co-stained with αPS1 integrin (green, left; inverted grayscale, right). Right panels display areas in the Anterior (A) and Posterior (P) compartments, which are outlined by rectangles. Scale bars: 50 μm (left); 10 μm (right). **E** Apical surface of the anterior (left) and posterior (right) compartments of discs expressing CD8-Cherry with diß in the posterior compartment. Cell outlines are visualized with E-cad-GFP (inverted grayscale). Scale bar: 5 μm. **F** The apical cell area of wing discs depicted in E, with each dot representing the cell average of an individual disc; compartments from same disc are connected by grey lines (n=22 discs). **G** Wing pouch region of the disc expressing CD8-Cherry with diß in the posterior compartment, visualised with E-cad-GFP (green, left), CD8 (magenta, left) and cleaved caspase-3 (Dcp-1, yellow, left; inverted grayscale, right and bottom sagittal projection of the dashed line. Scale bars: 50 μm (top) and 20 μm (bottom).

**Figure EV3.**
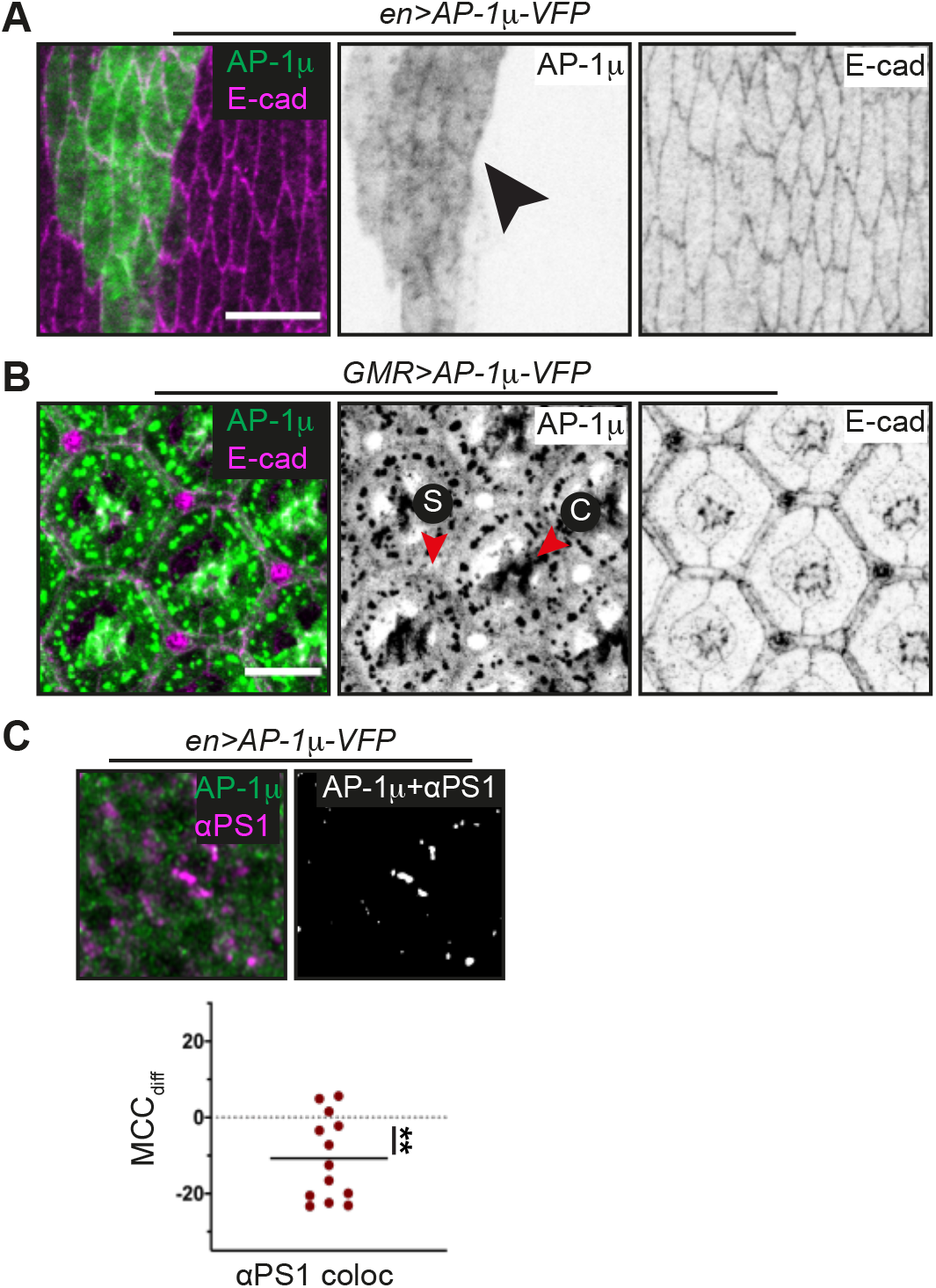
Characterization of the apical pool of AP-1 in *Drosophila* tissues. Related to Figure 2. **A** Apical view of stage 15 dorsal embryonic epidermis expressing AP-1μ-VFP (green, left; inverted grayscale, centre) in the posterior compartment and co-stained with E-cad (magenta, left; inverted grayscale, right). Arrowheads indicates cell borders. Scale bar: 10 μm. **B** Distal view of retina at 45% pupal development expressing AP-1μ-VFP (green, left; inverted grayscale, centre) and co-stained with E-cad (magenta, left; inverted grayscale, right). Arrowheads indicate AP-1μ-VFP at the AJs. Scale bar: 10 μm. **C** Example of colocalization (top, right) in the original image (left, extracted from Fig 2A) and the Mander’s Correlation Coefficient difference (MCC_dif_, see Methods) for the αPS1 and AP-1μ-VFP threshold signal (bottom, n=13). **P<0.01 (one sample t-test).

**Figure EV4.**
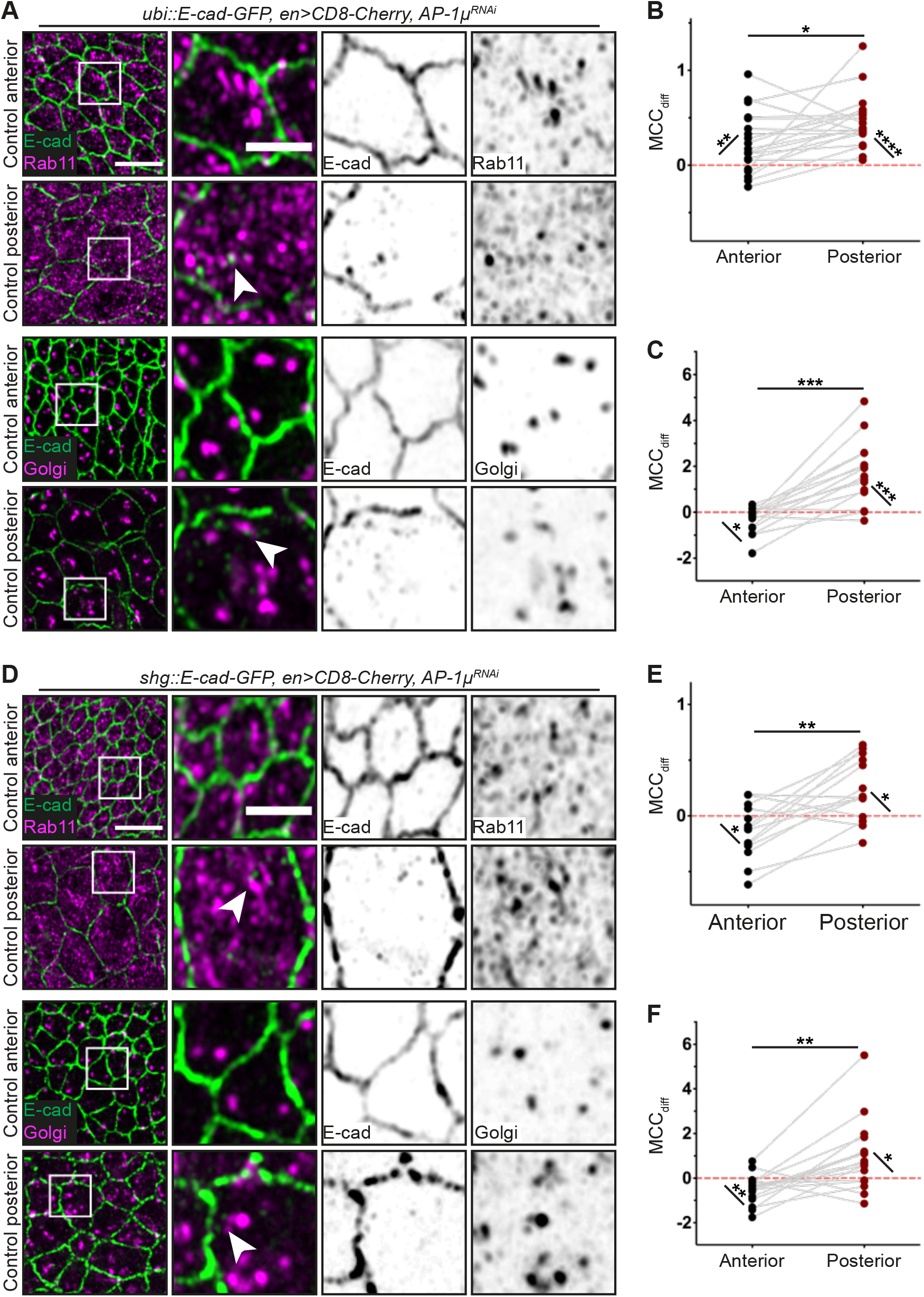
AP-1 knockdown increases the intracellular pool of E-cadherin. Related to Figure 3. **A** Dorsal wing pouch regions of discs co-expressing *ubi::E-cad-GFP* (green) with CD8-Cherry (not shown) and AP-1μ RNAi in the posterior compartment. Discs are stained for Rab11 (top two rows) and Golgi mab (bottom two rows), both depicted in magenta. Right panels show magnified regions within the white squares in left panels. Arrowheads indicate co-localization of E-cad and each marker. Scale bars: 5 μm (left) and 2 μm (detail). **B** Manders’ Correlation Coefficient difference (MCC_dif_, see Methods) for the co-localization between Rab11 and *ubi::E-cad-GFP.* Each dot represents average for individual paired compartments (n=20 discs). *P<0.05 (Wilcoxon test); **P<0.01 and ***P<0.001 (one sample t-test). **C** Manders’ Correlation Coefficient difference (MCCdiff, see Methods) for the co-localization between Golgi and *ubi::E-cad-GFP*. Each dot represents average for individual paired compartments (n=15 discs). ***P<0.05 (Wilcoxon test and one sample t-test); *P<0.01 and ***P<0.001 (one sample t-test). **D** Dorsal wing pouch regions of discs co-expressing *shg::E-cad-GFP* (green) with CD8-Cherry (not shown) and AP-1μ RNAi in the posterior compartment. Discs are stained for Rab11 (top two rows) and Golgi mab (bottom two rows), both depicted in magenta. Right panels show magnified regions within the white squares in left panels. Arrowheads indicate co-localization of E-cad and each marker. Scale bars: 5 μm (left) and 2 μm (detail). **E** Manders’ Correlation Coefficient difference (MCC_dif_, see Methods) for the co-localization between Rab11 and *shg::E-cad-GFP*. Each dot represents average for individual paired compartments (n=14 discs). *P<0.05 (one sample t-test) and **P<0.01 (Wilcoxon test). **F** Manders’ Correlation Coefficient difference (MCC_dif_, see Methods) for the co-localization between Golgi and *shg::E-cad-GFP*. Each dot represents average for individual paired compartments (n=17 discs). *P<0.05 (one sample t-test) and **P<0.01 (Wilcoxon test and one sample t-test).

